# Targeting Kinases to Reshape the Immunopeptidome landscape in Chronic Myeloid Leukemia

**DOI:** 10.1101/2025.10.07.680859

**Authors:** Veronica Venafra, Maria Wahle, Giorgia Massacci, Valeria Bica, Patrizia Chiusolo, Dimitrios Mougiakakos, Martin Boettcher, Thomas Fischer, Livia Perfetto, Matthias Mann, Francesca Sacco

## Abstract

Antileukemia immunity is essential for disease control and maintaining treatment-free remission in chronic myeloid leukemia (CML). Immunotherapeutic approaches hold significant promise for enhancing immune-mediated disease control. However, successful immunotherapy relies on identifying CML-associated antigens and achieving robust antigen presentation. Protein kinases are central regulators of signaling, and protein turnover, influencing which peptides are processed and presented on HLA molecules. Indeed, previous works with CDK4/6 or MEK inhibitors showed that kinases influence immune recognition of cancer cells. However, the molecular mechanisms underlying the modulation of the immunopeptidome by kinases inhibitors treatment remained unclear. Here, we investigated whether pharmacological inhibition of key kinases, including BCR-ABL, JNK and LCK, could be leveraged to reshape the immunopeptidome of CML cells, enhancing the exposure of specific tumor-associated antigens. By employing high-sensitive mass spectrometry (MS)–based immunopeptidomics, we quantified more than 20,000 HLA-I peptides. Pharmacological suppression of LCK, JNK and BCR-ABL drastically remodels the antigen landscape, impacting the exposure of about 4,000 HLA-I ligands. By integrating immunopeptidomics, phospho-immunopeptidomics, and proteomic, our study revealed three complementary molecular mechanisms of kinase-dependent antigen control: direct phosphorylation of HLA-I peptides (i), modulation of source protein stability (ii), and control of transcription factors governing antigen expression (iii). Moreover, comparative HLA ligandome analysis of benign hematological specimens and CML primary samples identified a panel of about 90 frequently presented CML-exclusive peptides, whose presentation can be significantly increased by LCK and JNK inhibition. Our study establishes a framework to rationally combine kinase inhibitors with immunotherapies to enhance antigen visibility and improve antileukemic immunity.

## Introduction

Chronic myeloid leukemia (CML) is defined by the chromosomal translocation t(9;22), which results in the BCR-ABL fusion gene ^1^. This fusion protein exhibits constitutive tyrosine kinase activity and is effectively targeted by approved tyrosine kinase inhibitors (TKIs), which have significantly improved the prognosis for CML patients ^2^. The current therapeutic objective in CML management is to achieve treatment-free remission (TFR) after discontinuation of BCR-ABL TKI treatment. TKI treatment is discontinued once stable and long-term complete or major molecular remission is achieved. However, only a subset of patients can stop TKI therapy without experiencing molecular relapse ^3^. As a result, long-term TKI therapy remains the standard of care for most patients, despite its potential for considerable side effects and the development of resistance ^4^. Multiple studies indicate that immunological control may play a crucial role in promoting TFR. Restoration of immune function - characterized by heightened activity of natural killer (NK) cells and T cells, along with decreased PD-1 expression on T cells ^5^ - has been linked to the successful achievement of TFR. Therefore, enhancing CML-specific immune responses through T cell–based immunotherapy offers a potential route to increase the proportion of patients achieving durable treat-free remission or even cure. Nonspecific immunotherapy strategies, including allogeneic stem cell transplantation and interferon-α (IFN-α) therapy, have been shown to induce long-lasting remissions following TKI discontinuation ^6^. Immune checkpoint inhibitors, which have transformed treatment paradigms for many solid tumors, are now under investigation in the context of CML ^7^.

More targeted immunotherapeutic approaches involve agents that induce immune responses against leukemia-specific antigens, such as peptide vaccines, T cell receptor (TCR)-mimic antibodies, and genetically engineered T cells ^8,9^. These strategies depend on the identification of tumor-associated, HLA-presented peptides that are recognized by T cells. While neoepitopes resulting from tumor-specific mutations are key targets in solid tumors with high mutational burdens, their role in cancers with lower mutational loads, like CML, remains uncertain. Advances in mass spectrometry (MS)-based immunopeptidomics have enabled large-scale identification of nonmutated, tumor-associated HLA peptides that can elicit peptide-specific T cell responses and serve as targets for T cell–based therapies ^10^. Importantly, successful immunotherapy relies on robust antigen presentation by cancer cells, which may be impaired by several factors, including aberrant signaling pathways. Pharmacological inhibition of specific kinases - such as CDK4, CDK6, and MEK - has been shown to enhance cancer cell immunogenicity by increasing tumor-associated antigen (TAA) presentation in other cancer types ^11,12^.

However, a comprehensive characterization of the crosstalk between signaling pathways and TAA exposure is still lacking. Gaining deeper insight into this complex interplay is critical, as numerous FDA-approved kinase inhibitors are already in clinical use and, in principle, could be leveraged to enhance the immunogenicity of cancer cells. To investigate this, we initially employed a computational network-based approach to identify kinases potentially involved in regulating antigen presentation. Subsequently, we applied high sensitive mass spectrometry (MS)-based proteomics to unbiasedly quantify alterations in class I HLA peptide repertoires following treatment with three different kinase inhibitors, including the standard-of-care therapy, imatinib. This analysis revealed enhanced presentation of selected HLA-I peptides, including novel TAAs, in response to specific kinase inhibitors.

To elucidate the signaling-driven mechanisms underlying these alterations, we combined our immunopeptidomic data with mass spectrometry–based (phospho)immunopeptidome and (phospho)proteome profiling of kinase inhibitor–treated cells and a computational network-driven approach. Through this integrative strategy, we identified for the first time three complementary phosphorylation-dependent molecular mechanisms that regulate extracellular antigen presentation: direct phosphorylation of HLA-I peptides, modulation of source protein stability, and control of transcription factors governing antigen expression. Our findings underscore selective kinase inhibition as a strategy to boost immunotherapy by enhancing the presentation of target antigens. Importantly, by characterizing the molecular mechanisms and signaling pathways that govern HLA-I antigen presentation, this work establishes a quantitative, network-based immunopeptidomics framework for systematically identifying and exploiting drug-induced HLA-I peptide exposure in next-generation immunotherapies.

## Results

### Identification of kinases controlling antigen presentation

To systematically identify key signaling molecules involved in regulating tumor-associated antigen (TAA) presentation, we applied the ProxPath algorithm, which estimates the functional proximity of signaling proteins to a target pathway using curated causal interactions captured in the SIGNOR resource ^13^. We focused on identifying druggable kinases, playing key roles in CML, that are functionally connected to the antigen processing and presentation machinery (APPM) pathway. This analysis highlighted three kinases - BCR-ABL1, JNK and LCK - as being linked to the APPM via causal paths associated with either upregulation or downregulation of antigen presentation (**Fig. S1A**). Among them, LCK showed the highest number of connections to the APPM (56 paths), suggesting a strong regulatory relationship.

### Immunopeptidomic profiling of kinase-inhibited CML cells

First, we investigated the impact of pharmacological inhibition of these three kinases on the antigen exposure. To this aim, BV173 cells were treated for 72h with imatinib (BCR-ABL inhibitor), SP600125 (JNK inhibitor) and PP2 (LCK inhibitor). Under these conditions, we employed the recently developed IMBAS (Immunopeptidomics by Biotinylated Antibodies and Streptavidin) workflow ^14^ in combination with high-sensitivity MS-based proteomics to quantify MHC-I-bound peptides in biological triplicates (**Fig. 1A**). Specifically, biotinylated W6/32 antibodies were used to immunoprecipitate antigenic peptides presented by MHC-I molecules, which were then captured on streptavidin beads. After washing, elution, and molecular-weight filtered, samples were loaded onto Evotips for downstream mass spectrometry analysis. Our approach enabled the identification of 21,078 unique HLA-I peptides (**Fig. S1B**) across the four experimental conditions. As expected, the peptides were highly enriched for nonameric sequences (**Fig. S1C**). Most of the peptides (20,081 out of 21,078) were predicted to be strong or weak binders of the allelic profile of BV173 cells, by using NetMHCpan 4.0 ^15^ (**Fig. 1B, Table S1**).

**Figure 1.**
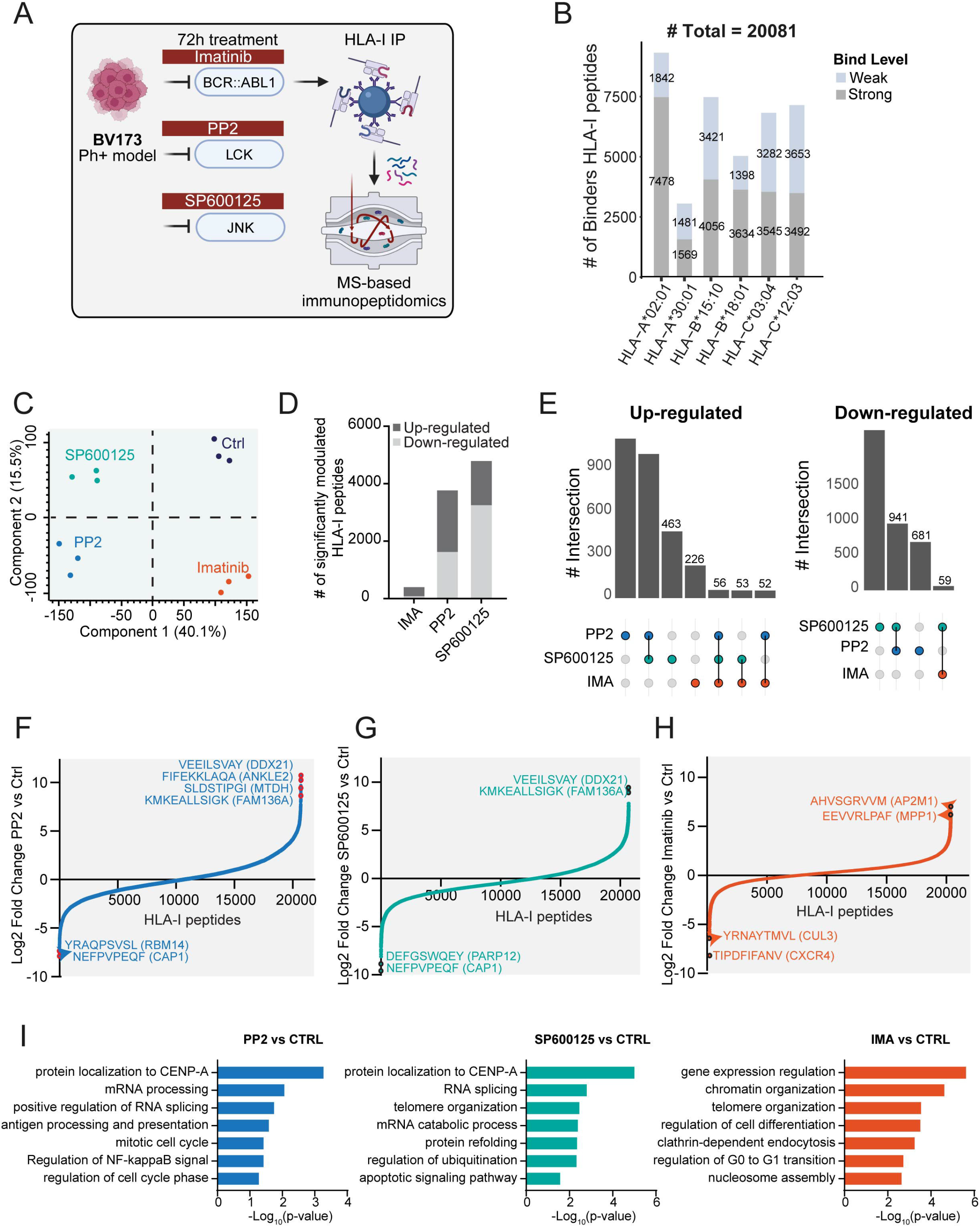
Quantitative analysis of HLA-I immunopeptidome profiling across kinase inhibitor treatments. **A.** Experimental workflow depicting the use of MS-based immunopeptidomics on the BV173 Ph+ chronic myeloid leukemia model treated for 72 hours with 3 kinase inhibitors. **B**. Number of peptides predicted by NetMHCpan 4.1 ^15^ to bind distinct BV173 HLA-I alleles, separated by strong (grey) and weak (skyblue) binders. **C**. Principal component analysis (PCA) of 21,078 HLA-I peptides quantified in BV173 cells treated with three different kinase inhibitor treatments (Imatinib, PP2, SP600125) and control conditions. **D**. Number of significantly up- and down-regulated HLA-I peptides for each kinase inhibitor treatment relative to control. Significance was assessed using two-sample t-tests (FDR < 0.05). **E**. Upset plots display the intersections of significantly upregulated (left) and downregulated (right) peptides among the different kinase inhibitor treatments. **F–H**. Waterfall plots showing log2 fold changes in HLA-I peptide abundance upon treatment with PP2 (**F**), SP600125 (**G**), and Imatinib (**H**) compared to control. Selected peptides with significant and strong fold changes and their source proteins are annotated.

To assess how different kinase inhibitors influence the immunopeptidome, we analyzed the global changes in HLA-I-bound peptides following targeted treatments. First, to investigate whether the drugs could be classified in an unsupervised manner based on their impact on HLA-I peptide presentation, we applied principal component analysis (PCA) to the immunopeptidomic dataset. We observed a strong reproducibility and distinct clustering of the treatments, which was further supported by unsupervised hierarchical clustering (**Fig. 1C, Fig. S1D**). These analyses revealed a direct and regulated link between kinase inhibition and antigen presentation. Interestingly, we observed a stronger similarity in the immunopeptidome response to LCK and JNK inhibition (PP2 and SP600125) compared to BCR-ABL inhibition with imatinib.

To identify HLA-I peptides significantly affected by kinase inhibitor treatments, we performed a stringent statistical t-test analysis (**Table S2**). SP600125 and PP2 modulated over 4,000 HLA-I peptides, whereas imatinib showed a milder effect, altering the presentation of approximately 200 peptides (**Fig. 1D**). SP600125 and PP2 both induced substantial up- and down-regulation of HLA-I peptides, with a notable overlap in the modulated peptides (**Fig. 1E**, up- and down-regulation intersections). Notably, treatments with PP2 and SP600125 induced stronger alterations in the abundance of surface-presented HLA-I peptides, with some antigens showing changes in expression spanning up to 3 orders of magnitude (**Fig. 1F–G**). Imatinib also reshaped the immunopeptidome, with HLA-I peptides being either upregulated or downregulated by as much as 64-fold (**Fig. 1H**). Importantly, the changes induced by SP600125, PP2, and imatinib were observed across all major HLA-I subtypes (HLA-A, -B, and -C), indicating that their effects are not restricted to specific HLA alleles (**Fig. S1E**). Consistently, sequence motif analysis of peptides significantly modulated by SP600125, PP2, and imatinib revealed strong conservation with known binding motifs of the HLA alleles expressed in BV173 cells, particularly at anchor positions (**Fig. S2A-C**), further validating the biological relevance of these changes. We also performed GO term enrichment analysis on the source proteins of the HLA-I peptides whose abundance is significantly upregulated by the three kinase inhibitors. Our analysis identified enriched biological processes of interest, including cell cycle, chromatin organization, cell division, and antigen presentation. These enrichments consistently reflect the expected biological response to imatinib, SP600125 and PP2 (**Fig. 1I**).

### Multi-omics elucidates kinase-dependent mechanisms shaping HLA-I exposure

Our findings show that inhibition of selected kinases leads to strong and selective up- or downregulation of hundreds of HLA-I peptides. To determine whether this effect is due to direct modulation of the antigen processing and presentation machinery, we assessed the MHC-I surface expression in BV173 cells after 72 hours of treatment with PP2, SP600125, and imatinib using flow cytometry. MHC-I expression levels remained unchanged across all treatments, suggesting that JNK, LCK, and BCR-ABL regulate antigen exposure through alternative mechanisms (**Fig. S2D**). As alterations in signaling, phosphorylation, and protein abundance are ultimately integrated into the repertoire of HLA-I–presented peptides, the immunopeptidome reflects the intracellular state. Thus, to dissect the molecular mechanisms underlying kinase-dependent modulation of HLA-I peptide presentation, we developed a network-based strategy integrating multi-omic datasets with the human naïve causal network (**Fig.□2A**). Specifically, in addition to immunopeptidomic profiling, we examined three additional regulatory layers in BV173 cells treated with kinase inhibitors: phospho-immunopeptidomics, global phosphoproteomics, and proteomics. First, we re-analyzed the immunopeptidomic dataset to specifically search for phosphorylated HLA-I peptides (phospho-immunopeptidomic dataset). Next, we employed high sensitive MS-based proteomics to quantify changes on the global proteome and phosphoproteome in BV173 upon 72h of treatment with PP2, SP600125 and imatinib. By this strategy, we quantified about 60 phosphorylated HLA-I peptides (**Table S3**), more than 7700 proteins (**Table S4 Fig. 2B**) and 17,000 phosphosites (**Table S5, Fig. 2B**). Principal component analyses of the proteomics and phosphoproteomics datasets show high reproducibility and distinct separation among treatments (**Fig. S3A-B**). Statistical analyses (FDR <0.05) revealed that 45 phospho-HLA-I peptides (**Table S3**), 4,400 phosphosites, and 4,700 proteins were significantly modulated by at least one kinase inhibitor (**Fig. 2C, Fig.S3C-D**). We then integrated these multi-omic datasets with the human naïve causal network to uncover the mechanisms by which kinase inhibition influences HLA-I peptide presentation. Specifically, we found that BCR-ABL, JNK, and LCK kinases reshape the CML immunopeptidome through three distinct regulatory mechanisms: (i) direct modulation of peptide phosphorylation, (ii) stability-dependent control of source proteins, and (iii) transcription factor–mediated regulation of peptide abundance. This approach allowed us to systematically map the relationships between kinase signaling, phosphoproteome dynamics, and HLA-I peptide presentation. Finally, we implemented a multi-step strategy to assign each modulated HLA-I peptide to one of the following regulatory modes, when possible.

**Figure 2.**
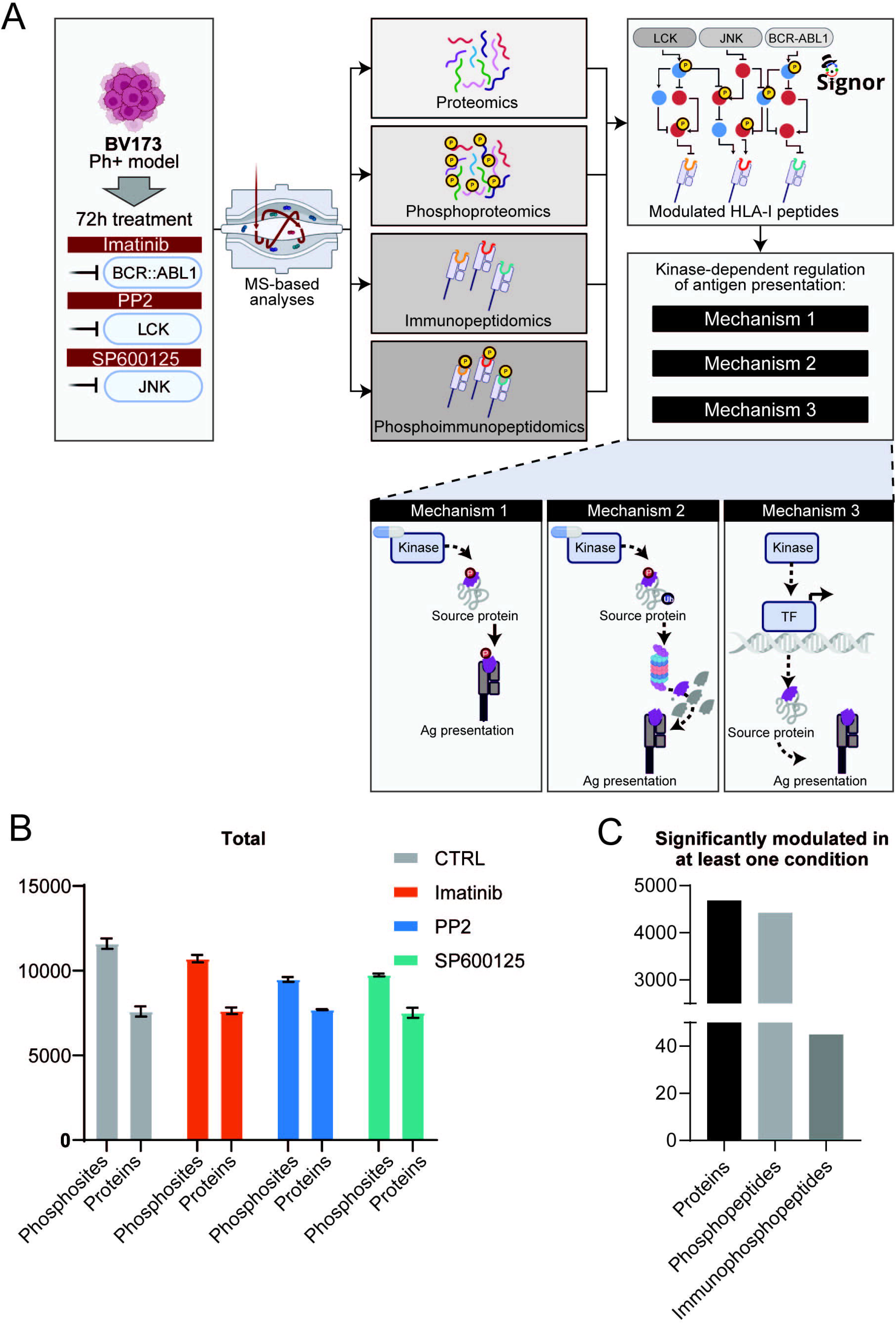
Application of a multi-step strategy to investigate treatment-dependent HLA-I modulation. **A.** Experimental workflow depicting the use of four different MS-based layers to ultimately identify the molecular mechanisms through which kinase inhibition can lead to a modulation of HLA-I expression. **B**. Total number of proteins and phosphopeptides identified across treatments and control. **C**. Number of significantly modulated proteins, phosphopeptides, and immunophosphopeptides in at least one condition compared to control.

### Kinase-dependent HLA-I phosphorylation controls antigen exposure

Phosphorylation of HLA-I peptides has been correlated with enhanced affinity for HLA-I molecules ^16^. To assess the extent to which kinase inhibitors reshape the immunopeptidome by directly influencing the phosphorylation of HLA-I peptides, we focused on the 45 modulated phospho-HLA-I peptides and used the SIGNOR database to identify those that (i) contained a phosphosite with a known upstream kinase and (ii) whose upstream kinase is itself directly or indirectly regulated by the inhibitor treatment (**Fig. 3A**). When available, we used our phosphoproteomic dataset (**Table S5**) to assess the activity of the upstream kinases upon drug treatment. Even if limited in number, direct phosphorylation-dependent regulation of HLA-I peptides was confirmed. Specifically, three HLA-I peptides were found to be indirectly phosphorylated by BCR-ABL, JNK and LCK kinases: FSVASPLTL, KNITPRKK, and VKNITPRKK (Table S6). Among these, inhibition of JNK and BCR-ABL significantly reduced the presentation of the phosphorylated SCML2-derived HLA-I peptide KNITPRKK (**Fig. 3B**). Network analysis revealed that both imatinib and PP2 treatments led to inactivation of CDK1/2 activity, as shown by decreased phosphorylation at the T161 activation site. This loss of CDK1/2 activity resulted in decreased phosphorylation and surface presentation of the SCML2-derived HLA-I peptide (KNITPRKK), which contains the CDK1/2 substrate residue T305. Our findings indicate that kinase inhibitors can modulate HLA-I peptide presentation through phosphorylation-dependent mechanisms, where drug-induced inactivation of upstream kinases alters substrate phosphorylation and consequently the repertoire of peptides displayed at the cell surface.

**Figure 3.**
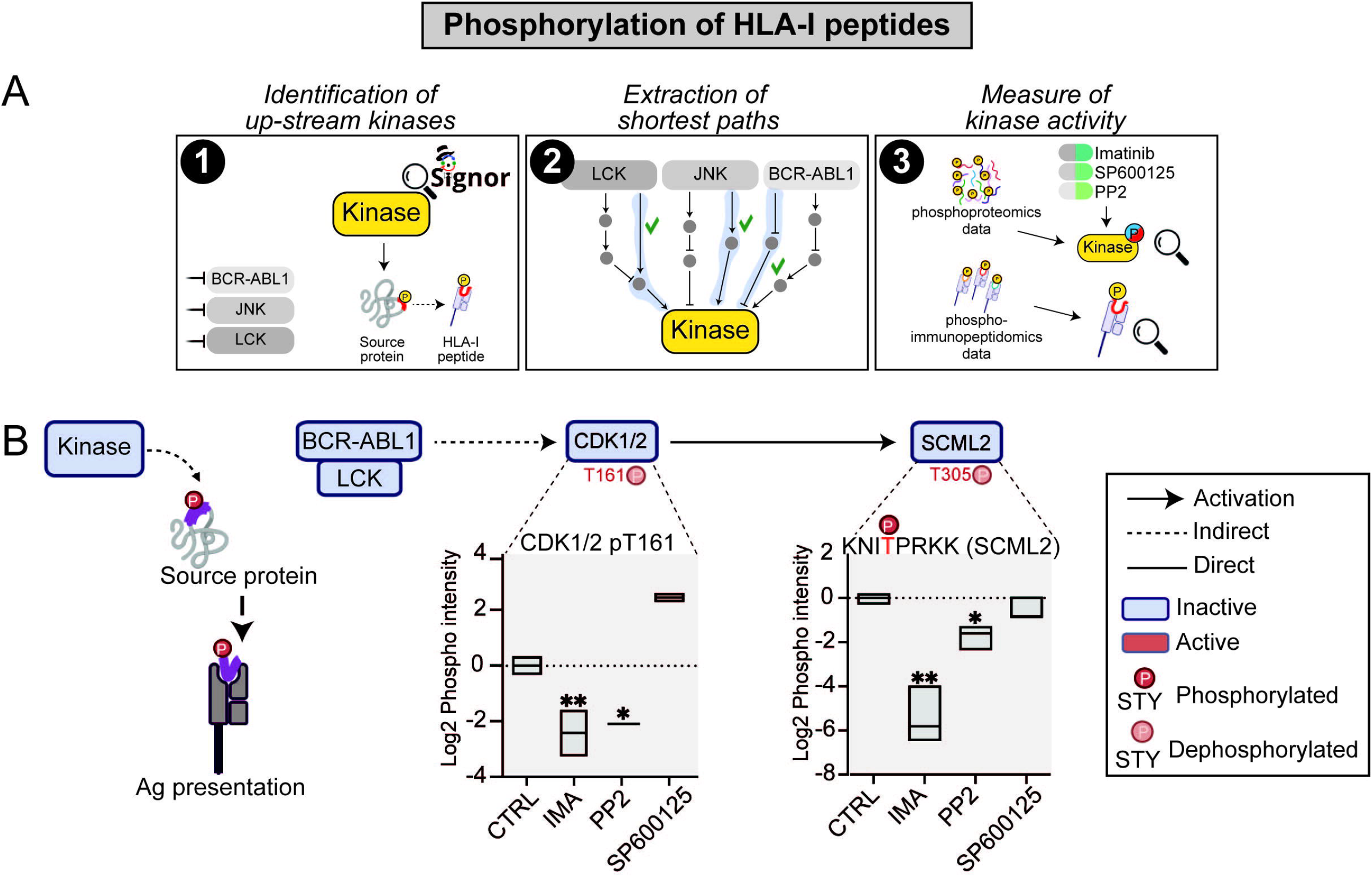
Phosphorylation-dependent modulation of HLA-I peptides. **A.** Illustration of the multi-step strategy we applied to identify HLA-I peptides modulated via direct phosphorylation. **B**. Representative example illustrating how BCR-ABL and LCK kinases influence HLA-I peptide phosphorylation. Boxplots show the log2 phospho-intensity of the activating site T161 on CDK1/2 (from phosphoproteomic data) and the log2 phospho-intensity of the SCML2-derived HLA-I peptide (from phospho-immunopeptidomic data).

### Kinase-mediated control of source-protein stability modulates HLA-I exposure

Phosphorylation of key residues can impact the proteasome-mediated degradation of proteins and consequently the HLA-I peptide exposure. We selected HLA-I peptides that were significantly modulated by at least one kinase inhibitor and derived from source proteins with phosphosites known to regulate stability (**Table S2**), as annotated in the SIGNOR database (**Fig. 4A**). We further prioritized those whose upstream kinases were modulated by the inhibitors. When available, we used our phosphoproteomic dataset to assess the activity of the upstream kinases and the phosphorylation level of the antigen source proteins upon kinase inhibition. This phosphorylation-dependent modulation of protein stability emerged as a more prevalent mechanism, involving about 20 HLA-I peptides (**Table S6**). Interestingly, we found that JNK inhibition leads to a significant decrease in the presentation of two Vimentin (VIM)-derived HLA-I peptides. Mechanistically, our network-based strategy revealed the mechanism underlying the JNK-dependent modulation of these VIM-derived HLA-I peptides. Specifically, JNK inhibition leads to AKT inactivation by reducing phosphorylation at its activation site T450, as revealed by our phosphoproteomic analysis. In its inactive state, AKT no longer phosphorylates VIM at S39, as confirmed by our phosphoproteomic data (Table S5). This results in increased VIM stability and decreased presentation of VIM-derived HLA-I peptides (**Fig. 4B**). These results demonstrate that kinase-dependent phosphorylation can regulate HLA-I peptide presentation by modulating source protein stability, with changes in protein degradation, and ultimately the repertoire of peptides displayed on the cell surface.

**Figure 4.**
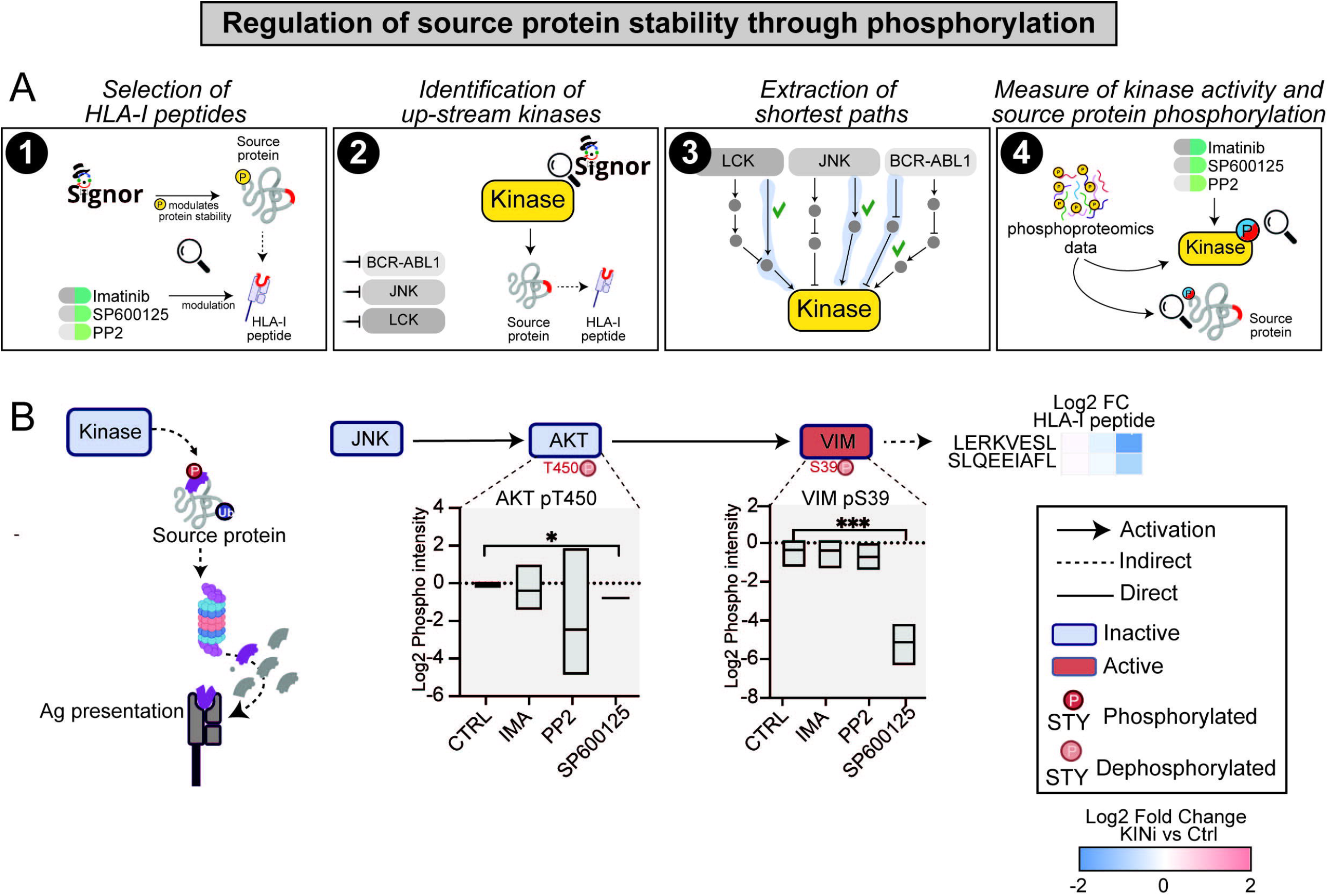
Regulation of source protein stability. **A.** Illustration of the multi-step strategy we applied to identify HLA-I peptides modulated via regulation of source protein stability. **B**. Model depicting how JNK inhibition regulates VIM stability via phosphorylation of key regulatory residues. Boxplots display phosphorylation levels of AKT (pS450) and VIM (pS39) following inhibitor treatment (phosphoproteomic data). The heatmap summarizes the log2 fold changes in abundance of VIM-derived peptides after kinase inhibition.

### Kinase regulation of source-protein transcription impacts HLA-I exposure

Phosphorylation also affects transcription factor (TF) activity, thereby influencing gene expression and, consequently, antigen presentation. We identified HLA-I peptides derived from source proteins transcriptionally regulated by transcription factors whose activity is modulated by targeted kinases (**Fig**.□**5A**). When the data were available, our phosphoproteomic dataset was used to assess the phosphorylation of transcription factors, while our proteomic and immunopeptidomic datasets were leveraged to monitor the abundance of the downstream target source proteins and HLA-peptides, respectively. We found that the transcriptional regulation represented a major layer of control, with 16 transcription factors identified as regulators of 170 source proteins and 222 associated HLA-I peptides (Table S6). For instance, our network-based strategy identified STAT3 and MYC as central nodes, collectively modulating ∼20 source proteins and their corresponding HLA-I peptides. Analysis of our (phospho)proteomic datasets revealed that inhibition of BCR-ABL, JNK, and LCK significantly reduced phosphorylation of STAT3 at S727 and MYC at T73, their respective activating residues. This reduction in phosphorylation was associated with a marked decrease in the protein abundance of STAT3- and MYC-regulated source proteins and in the presentation of their derived HLA-I peptides, as shown in the heatmaps (**Fig. 5B**). These findings highlight transcriptional regulation as a major mechanism by which kinase signaling shapes HLA-I peptide presentation. Kinase-dependent phosphorylation of transcription factors can modulate the abundance of their downstream proteins and the surface-presented peptide repertoire.

**Figure 5.**
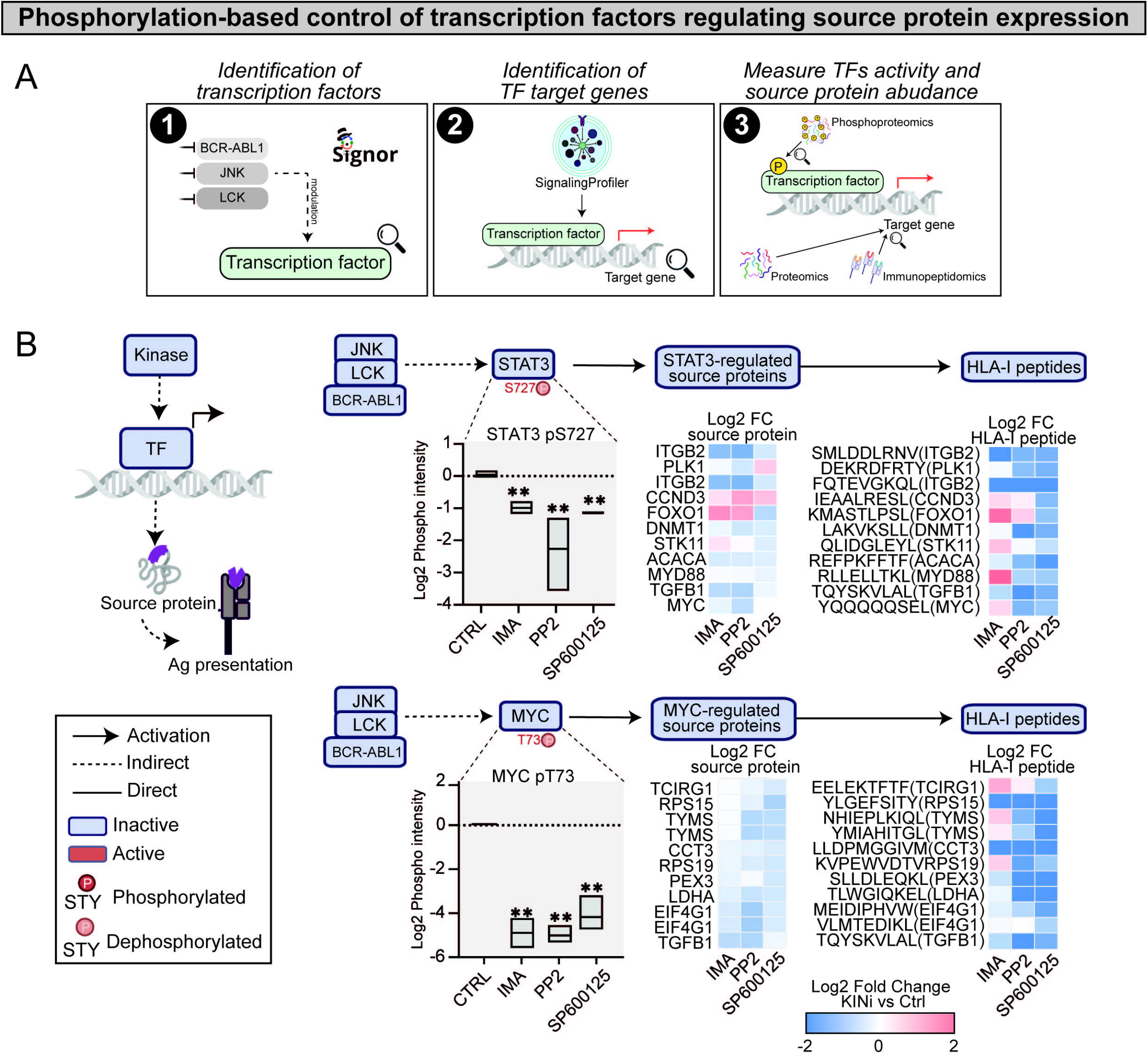
Transcriptional regulation of source proteins. **A.** Illustration of the multi-step strategy we applied to identify HLA-I peptides modulated via transcriptional regulation of source protein. **B**. Model showing how kinase inhibitors modulate transcription factors that control the expression of antigen source proteins. Boxplots present phosphorylation levels of STAT3 (pS727) and MYC (pT73) in response to inhibitor treatment (phosphoproteomic data). Heatmaps illustrate log2 fold changes in the abundance of STAT3- and MYC-regulated source proteins (proteomic data) and their corresponding HLA-I peptides (immunopeptidomic data).

### LCK, BCR-ABL and JNK kinases modulates the exposure of specific TAAs

Our data show that inhibition of LCK, JNK and, to a lesser extent BCR-ABL, significantly reshapes the immunopeptidome profile of a CML cell line. We next asked whether kinase inhibition may also impact the abundance of well characterized tumour associated antigens (TAAs). To address this question, we systematically compared our cell line–derived immunopeptidomic dataset with two independent reference cohorts: the immunopeptidome of a cohort of 21 primary CML samples and a comprehensive ligandome of benign hematologic tissues ^10^ (**Fig. 6A**). This integrative analysis revealed that approximately 10% of the HLA-I peptides identified in our study were also detected in primary CML samples, but were entirely absent from the atlas of benign hematologic tissues (**Fig. 6B**). These 255 peptides were classified as TAAs and were found to be presented in about 10% of CML patients, highlighting their potential significance as immunotherapy targets (**Fig. 6C**). In addition to identifying novel tumor-associated antigens (TAAs), we also assessed whether our dataset of naturally presented HLA-I peptides included well-characterized cancer-testis antigens (CTAs) ^17,18^. CTAs are normally restricted to germ line cells (in testis and placenta) but are aberrantly over expressed in several tumors, representing promising immunotherapy targets ^19^. Our analysis identified 17 distinct HLA class I peptides derived from 15 CTAs. Notably, 7 out 17 antigens were also detected in benign hematologic samples (**Fig. 6D**) and only one antigen was identified in both CML and healthy primary samples. Consistent with this observation, frequent tumor-exclusive presentation of CTAs has not been observed in a large-scale immunopeptidome profiling of 21 CML patients ^10^. This underscores the relevance of identifying perturbations enhancing the exposure of these 255 CML-exclusive, non-mutated antigens representing a unique antigenic repertoire whose presentation may be therapeutically exploitable. Thus, we evaluated how treatment with imatinib, PP2, and SP600125 influenced the abundance of these 255 TAAs. Quantitative immunopeptidomic profiling showed that imatinib had only a modest impact on TAA presentation. By contrast, PP2 and SP600125 treatment substantially altered the exposure of 52 and 65 TAAs, respectively, inducing strong up- or downregulation with fold changes ranging from fourfold to 64-fold (**Fig. 6E**). Remarkably, approximately 50 TAAs were predicted to have high immunogenicity and were significantly upregulated by at least one kinase inhibitor treatment (**Fig. 6F**).

**Figure 6.**
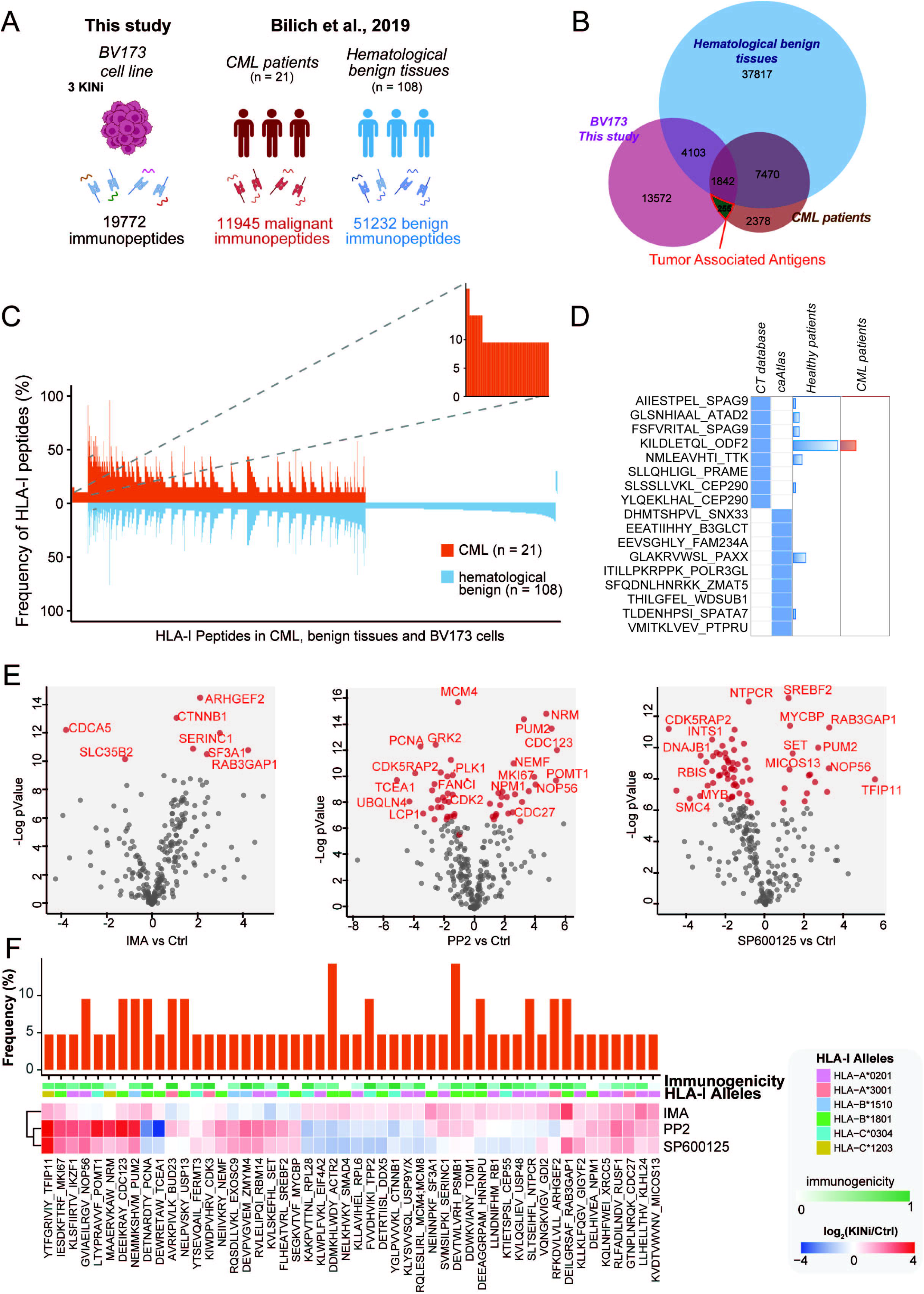
Identification and functional characterization of Tumor Associated Antigens. **A.** Cartoon reporting the comparison between this study’s dataset with the Bilich et al. (2019) immunopeptidome dataset. **B**. Venn Diagram reporting the overlaps in HLA-I peptide identification across BV173 binders, benign hematological tissues, and CML patient samples from Bilich et al., study. **C**. Frequency (y-axis) distribution of BV173 HLA-I peptides (x-axis) in CML patients and benign tissues ^20^. To allow for better readability, HLA peptides identified on >5% of the samples within the respective cohort are not depicted. The box on the left and its magnification highlight the subset of CML-associated antigens in BC4173 cell lines showing CML-exclusive high frequent presentation. **D**. Heatmap showing the cancer testis antigens, annotated in two databases ^17,18^ and identified in our study. Bar plots show the number of CML or healthy (hematological tissues) displaying the selected antigen. **E**. Volcano plots showing how kinase inhibition impacts the exposure of novel TAAs. **F**. Heatmap showing the log2 fold-change of 22 Tumor Associated Antigens significantly modulated in presentation by at least one kinase inhibitor and identified exclusively in CML patients. For each peptide, frequency in Bilich et al., CML patients (barplot), the DeepImmuno predicted immunogenicity (green color scale) and the HLA allele in BV173 cells with strongest affinity is reported.

## Discussion

In this study, we provide a comprehensive, multi-omic characterization of the signaling-driven mechanisms that regulate antigen presentation in CML cells. By integrating (phospho)-immunopeptidomics, global phosphoproteomics, proteomics, and a network-based computational framework, we demonstrate that inhibition of BCR-ABL, JNK, and LCK kinases profoundly reshapes the immunopeptidome of CML cells. Particularly, we show that pharmacological inhibition of kinase-specific signaling pathways not only alters the global abundance of HLA class I peptides but also selectively enhances the presentation of tumor-associated antigens (TAAs) with high predicted immunogenicity. These findings reveal novel opportunities for exploiting kinase inhibitors as adjuvants to immunotherapy in CML.

Consistent with this, MEK and CDK4/6 inhibition has previously been shown to sensitize tumors to immunotherapy by upregulating MHC class I molecules, promoting immune cell infiltration, activating T cells, and facilitating antigen recognition ^11,12^. While these previous studies have described a more general upregulation of antigen presentation upon kinase inhibition, our findings uncover a selective and fine-tuned mechanism of antigen modulation, highlighting a higher level of specificity in the rewiring of the immunopeptidome. Importantly, a key conceptual advance of our work is the identification of three distinct but complementary molecular mechanisms by which kinase inhibition controls antigen exposure. First, we show that direct phosphorylation of HLA-I peptides, although relatively rare, can be dynamically regulated by kinase signaling. Second, phosphorylation-dependent modulation of protein stability emerges as a more frequent mechanism, exemplified by the regulation of Vimentin-derived peptides via AKT signaling. Finally, we demonstrate that kinase inhibition profoundly impacts the transcriptional regulation of antigen source proteins, impacting the immunopeptidome landscape.

Our analysis uncovered a unique repertoire of 255 non-mutated TAAs that were consistently detected in primary CML samples but absent from benign hematologic tissues. This TAAs set represents a valuable resource for the design of immunotherapeutic strategies. Notably, classical cancer-testis antigens (CTAs) identified in our dataset, are also frequently detected in healthy hematopoietic samples. Such lack of CML exclusivity is coherent with previous large-scale immunopeptidomic studies, highlighting their limited clinical utility. By contrast, the 255 CML-exclusive peptides may provide more selective and safe targets for T cell–based therapies.

Together, these findings showcase how perturbation of kinase activity can extend across multiple molecular layers, ultimately reshaping the antigenic landscape. An important translational implication of our findings is that kinase inhibition can be leveraged to enhance the exposure of immunogenic antigens. While imatinib had only modest effects on TAA presentation, LCK and JNK inhibition caused extensive remodeling of the immunopeptidome, altering the abundance of more than 100 TAAs. Importantly, a subset of approximately 50 TAAs was highly immunogenic and also upregulated by kinase inhibitor treatment. Overall, these results suggest that rational combinations of kinase inhibitors with T cell–based immunotherapy may enhance antigen visibility and immune recognition, thereby increasing the likelihood of durable treatment-free remission in CML.

While these results are promising, some limitations of our study should be acknowledged. Although the 255 TAAs identified in our study were also presented in primary CML samples ^10^, further validation is needed to functionally confirm their immunogenicity and their ability to trigger effective anti-tumor T cell responses. Additionally, kinase inhibitors often display off-target effects, and their pleiotropic nature should be carefully considered.

In conclusion, our findings indicate that pharmacological inhibition of signaling molecules can serve as a strategy not only to kill cancer cells but also to enhance their immunogenicity by increasing tumor antigen visibility. To our knowledge, this is the first study that not only demonstrates that inhibition of specific kinases affects the presentation of specific HLA-I peptides, including novel TAAs, but also provides a mechanistic explanation for such kinase-dependent modulation. By delineating signaling-driven mechanisms and uncovering a set of CML-exclusive TAAs, we establish a framework for exploiting kinase inhibitors as tools to reshape the immunopeptidome and enhance the effectiveness of immunotherapy. Future studies will focus on validating the immunogenicity of these peptides and test rational drug–immunotherapy combinations in preclinical and clinical settings, with the ultimate goal of broadening the fraction of patients who achieve durable treatment-free remission.

## Authors’ contributions

Conceptualization, G.M., L.P., F.S.; resources, M.W., G.M., V.B., M.W.; software, V.V., L.P.; methodology, V.V., M.W., G.M., V.B., M.M., L.P., F.S.; formal analysis, V.V., M.W., G.M., V.B.; investigation, V.V., M.W., G.M., V.B, with the contribution of T.F., M.B., D.M.; writing original draft preparation, V.V., M.W., G.M., M.M., F.S.; writing review and editing, all; supervision, M.M., L.P., F.S.; funding acquisition, F.S. All authors have read and agreed to the published version of the manuscript.

## Acknowledgments

This research was funded by the Italian Association for Cancer Research (AIRC) with a grant to L.P. (MFAG Grant n. 28858) and a grant to F.S. (Start-Up Grant n. 21815) and by MUR PRIN 2022 (n. E53D23004850006). L.P. and F.S. are supported by a joint PRIN 2022 PNRR grant (n. P2022JRETW), funded by the European Union – NextGenerationEU, and by a SEED Sapienza Grant. G.M. is supported by MUR PRIN 2022 (n. 2022L8RAKN).

## Materials and Methods

### Cell culture

BV173 cell line were cultured in DMEM medium (Thermo Scientific, 41965062) supplemented with 10% heat-inactivated fetal bovine serum (Aurogene, AU-S1810-500), and 100 U/ml penicillin and 100 mg/ml streptomycin (Gibco 15140122). BV173 cells were treated at 500.000 cells/mL with the following inhibitors: imatinib (Selleck chemicals, S2475), PP2 (Selleck chemicals, S7008), and SP600125 (Selleck chemicals, S1460) for 72 hours.

### ProxPath analysis

The ProxPath algorithm was used to calculate the number of paths connecting each SIGNOR protein (input node) to the members of the *MHC class I antigen presentation* pathway (https://signor.uniroma2.it/pathway_browser.php?beta=3.0&organism=&pathway_list=SIGNOR-MCIAP&x=14&y=10). Input nodes with high number of paths impacting onto the antigen processing and presentation machinery (APPM) pathway, short distance with respect to the average distance (z-score < 0), and available inhibitors were selected for perturbation, namely LCK, BCR-ABL1 and JNK.

### Proteomic sample preparation

Cell pellets were dissolved in 100 µl lysis buffer (10% AcN, 60 mM TEAB, 5 mM TCEP, 25 mM CAA) and incubated at 76°C for 20 minutes on an orbital shaker. Next, samples were sonicated in a Bioruptor (Diagenode) for 10 cycles with 50% duty cycle. 2 µg of each trypsin and Lys-C were added for protein digestion and incubated overnight at 37°C. The digestion was stopped by adding formic acid (FA) to a final concentration of 1%. To clear them from cellular debris, samples were centrifuged at 5000 xg for 3 minutes in a benchtop centrifuge (Eppendorf). 5 µl of the digest were saved for full proteome measurement.

From the remaining digest, phopshorylated peptides were enriched on an AssayMAP Bravo Sample Prep Platform (Agilent), using the Phosphopetide Enrichment v2.1 Protocol in the Protein Sample Prep Workbench v3.2.0 with standard settings. 5 μl Fe-NTA Cartridges (Agilent) were primed with 0.1% trifluoroacetic acid (TFA) in AcN. Samples were loaded on the cartridges, washed with 0.1% TFA in 80% AcN and finally eluted in 0.5 M ammonium dihydrogenphosphate solution.

Both full proteome and phosphoproteome samples were loaded on Evotips (Evosep) according to the manufacturers protocol. In brief, Evotips were activated with isopropanol, primed two times with 50 µl of Buffer B (0.1% FA in AcN) and equilibrated two times with 50 μl of Buffer A (0.1% FA).

Samples were loaded on activated tips, washed twice with Buffer A and stored at 4°C until measurement. All centrifugation steps were done at 700 xg for 1 minute.

### Immunopeptidomics enrichment

Immunopeptidomics enrichment was performed using the IMBAS-MS workflow(ref). In brief, cell pellets were resuspended in 200µl lysis buffer (50 mM Tris, pH8;150 mM NaCl, 0.5% NP-40; 60 mM Octyl-b-D-glucopyranosid; 1 x cOmplete Protease inhibitor) and incubated for 30 min on ice. Samples were centrifuged for 20 min at 4°C at 15.000 rcf and the supernatant is transferred to a fresh tube. Samples were incubated over night with 10 µg W6/32 antibody at 4°C. Enrichment was performed using 20µl magnetic streptavidin beads. Beads were washed in 3 steps (1: 10 mM Tris pH8, 150 mM NaCl; 2: 10 mM Tris pH8, 450 mM NaCl; 3: 10 mM Tris pH8) before elution in 200 mM Glycine at pH2. The eluent was centrifuged through a 10kDa flat bottom tube filter (Merck) before loaded onto Evotips following the same procedure described above.

### Mass spectrometry analysis

The proteome and phosphoproteome LC/MS measurements were carried out using an EvoSep One liquid chromatography system (Evosep) coupled to a trapped ion mobility spectrometry quadrupole time-of-flight mass spectrometer (timsTOF HT, Bruker Daltonik) via a nanoelectrospray ion source. Samples were analyzed using a 44□min gradient (30 SPD) eluting the peptides at 250□nl/min flow rate. We used a 15□cm□×□75□μm column with 1.9 μm C18 beads (EvoSep) and a 10□µm ID zero dead volume electrospray emitter (Bruker Daltonik). Mobile phases A and B were 0.1% FA in water and 0.1% FA in AcN, respectively. For proteome measurements, 20 isolation windows were placed between an m/z range of 350 to 1200 and an ion mobility (IM) range between 1.3 and 0.7 V cm-2 at a 2.2 s cycle time. The collision energy was decreased as a function of the IM from 59 eV at 1/K0=1.6 to 20 eV at 1/K0 = 0.6. For phosphoproteome measurements the mass range was shifted to m/z 400 to 1400 and the ion mobility (IM) range was shifted to 1.45 and 0.75 V cm-2. The collision energy was adjusted towards 60 eV at 1/K0=1.5 to 54 eV at 1/K0=1.17 to 25 eV at 1/K0=0.85 and end at 20 eV at 1/K0=0.6.

The immunopeptidome LC/MS measurements were carried out using an EvoSep One LC system coupled to a timsTOF Ultra 2. Samples were separated using a 31 min gradient (whisper 40) over an Aurora IonOpticks 15 cm column. For acquisition, the same methods were employed as previously published (ref IMBAS paper).

### Proteome and phosphoproteome raw data processing

Full proteome raw files were analyzed with DIANN v1.9.2 and phosphoproteome samples with Spectronaut version 20. MS/MS spectra were matched against the Homo sapiens UniProtKB

FASTA database, with an FDR of < 1% at protein-, peptide- and modification-level. Enzyme specificity was set to trypsin with the maximum number of missed cleavages set to 1. Cysteine carbamidomethylation was added as a fixed modification, variable modifications were set to N-terminal protein acetylation and oxidation of methionine as well as phosphorylation of serine, threonine tyrosine residue (STY) for the phosphoproteomic samples.

### Immunopeptidome raw data processing

Immunopeptidome raw files were analyzed using DIA-NN v.2.0 against a predicted panHLA library ^14^. Results were filtered at a peptide FDR of 1% without applying a protein FDR. Phosphoimmunopeptidome results were obtained with Spectronaut version 20. Raw files were searched using the phosphor PTM workflow while setting the digestion type to unspecific and restricting the peptide length to minimum 7 and maximum 15. The peptide FDR was kept at 1% while the protein FDR filter was disabled. The PTM probability cut-off was set at 0.75.

### Data preprocessing

Immunopeptides’ source proteins were annotated using the HLA Ligand Atlas dataset ^20^ using immunorelated tissues (“Bone marrow”, “Lymph node”, “Myelon”, “Spleen”, “Thymus”) and the UNIPROT database. Intensities were log2 transformed, and only immunopeptides detected in 2 out of 3 replicates of at least one condition were retained. Missing values were imputed using the MinProb method (q = 0.01) from the DEP R package (v. 1.28.0) to simulate low-abundance signals.

### Differential expression analysis

Differentially modulated peptides between each kinase inhibitor and control were identified using Student’s t-test, with significance defined by an absolute fold-change > 0.5 and an adjusted p-value (Benjamini-Hochberg correction) < 0.05 calculated per sample.

### MHC binding affinity and immunogenicity prediction

Binding affinities of immunopeptides to HLA-I alleles were predicted using the NetMHCpan 4.1 web application ^15^, with the BV173 cell line HLA-I allele set (HLA-A02:01, HLA-A30:01, HLA-B15:10, HLA-B18:01, HLA-C03:04, and HLA-C12:03). Immunogenicity was assessed using the DeepImmuno web application (https://doi.org/10.1093/bib/bbab160), considering only peptides of length 9 or 10.

### Identification of Tumor-Associated Antigens

Tumor-associated antigens (TAAs) were defined as BV173 MHC-I-binding immunopeptides that were also detected in 21 CML patients from the Bilich et al., study ^10^ but not present in the benign peptide set reported in the same study. This resulted in a list of 255 peptides. From these, we selected as putative TAAs 94 immunopeptides that were significantly overexpressed in response to at least one kinase inhibitor.

### Mechanisms of regulations characterization

To characterize the mechanisms underlying the modulation of HLA-I peptide presentation, we integrated our multi-omic dataset with a literature-derived causal network.

Specifically, to investigate how kinase inhibition directly affects the exposure of phosphorylated HLA-I peptides (*Mechanism 1*), we applied the following strategy:

1. First, we queried the SIGNOR database to identify the upstream kinase of the phosphorylated HLA-I peptide source protein modulated upon kinase inhibition (**Table S2**).
2. Next, we extracted the shortest paths linking each upstream kinase to the inhibited kinases (BCR-ABL, LCK, JNK).
3. We then used our phosphoproteomic data (**Table S5**) to assess the activity of the upstream kinases upon drug treatment.

To identify HLA-I peptides modulated via phosphorylation-dependent changes in source protein stability (*Mechanism 2*), we used the following approach:

1. First, we selected only those HLA-I peptides significantly affected by kinase inhibition (**Table S2**) and mapped to source proteins whose phosphorylation has been reported to regulate protein stability in the SIGNOR database.
2. Next, we queried the SIGNOR database to identify the upstream kinase phosphorylating the source proteins.
3. We extracted the shortest paths linking each upstream kinase to the inhibited kinases (BCR-ABL, LCK, JNK), retaining only paths involving no more than two steps.
4. When the data were available, we used our phosphoproteomic data to assess the activity of the upstream kinases and the phosphorylation level of the source proteins upon kinase inhibition.

To identify HLA-I peptides modulated via phosphorylation-dependent modulation of transcription factors (*Mechanism 3*), we used the following strategy:

1. We used SIGNOR to identify transcription factors directly or indirectly regulated by the inhibited kinases.
2. We retrieved each transcription factor’s downstream target genes from the SignalingProfiler TF–target database (SIGNOR + Collectri).
3. When the data were available, phosphorylated transcription factors were compared with our phosphoproteomic data, while their downstream target genes were matched against the proteomic and immunopeptidomic datasets.
4. We retained HLA-I peptides showing consistent changes in expression aligned with both transcription factor activity and kinase inhibition.

### Flow cytometry analysis of MHC class I expression

BV173 cells were plated at a density of 1 × 10^6^ cells/mL and treated with each inhibitor (or DMSO) for 72h. After treatment, cells were washed with FACS buffer (1X PBS, 1 mM EDTA, 1% FBS, 0.1% NaN_3_) and incubated with anti-HLA Class I Alexa Fluor 647 conjugated antibody (R&D Systems™) using 1 μg antibody per 100 μL of cell suspension for 30 min at 4°C. Cells were washed 2X with FACS buffer, prior to analysis on CytoFLEX flow cytometer (Beckman Coulter Life Sciences). CytExpert software was used for data analysis. Cells were gated for live populations only. The median fluorescence intensity (MFI) was extracted and plotted for each sample as the average MFI +/– SD.

## Authors’ contributions

Conceptualization, G.M., L.P., F.S.; resources, M.W., G.M., V.B.; software, V.V., L.P.; methodology, V.V., M.W., G.M., V.B., L.P., F.S.; formal analysis, V.V., M.W., G.M., V.B.; investigation, V.V., M.W., G.M. V.B., with the contribution of T.F., M.B., D.M.; writing original draft preparation, V.V., M.W., G.M., L.P., F.S.; writing review and editing, all; supervision, L.P., F.S.; funding acquisition, F.S. All authors have read and agreed to the published version of the manuscript.

## Acknowledgments

This research was funded by the Italian Association for Cancer Research (AIRC) with a grant to L.P. (MFAG Grant n. 28858) and a grant to F.S. (Start-Up Grant n. 21815) and by MUR PRIN 2022 (n. E53D23004850006). G.M. is supported by MUR PRIN 2022 (n. E53D23004850006). L.P. and F.S. are supported by a joint PRIN 2022 PNRR grant (n. P2022JRETW), funded by the European Union – NextGenerationEU, and by a SEED Sapienza Grant. V.V. is supported by PON-MUR fellowship (n. DOT13IEP1U-1).

