## Supplementary figures S1-S3 for "Targeting Kinases to Reshape the Immunopeptidome landscape in Chronic Myeloid Leukemia"

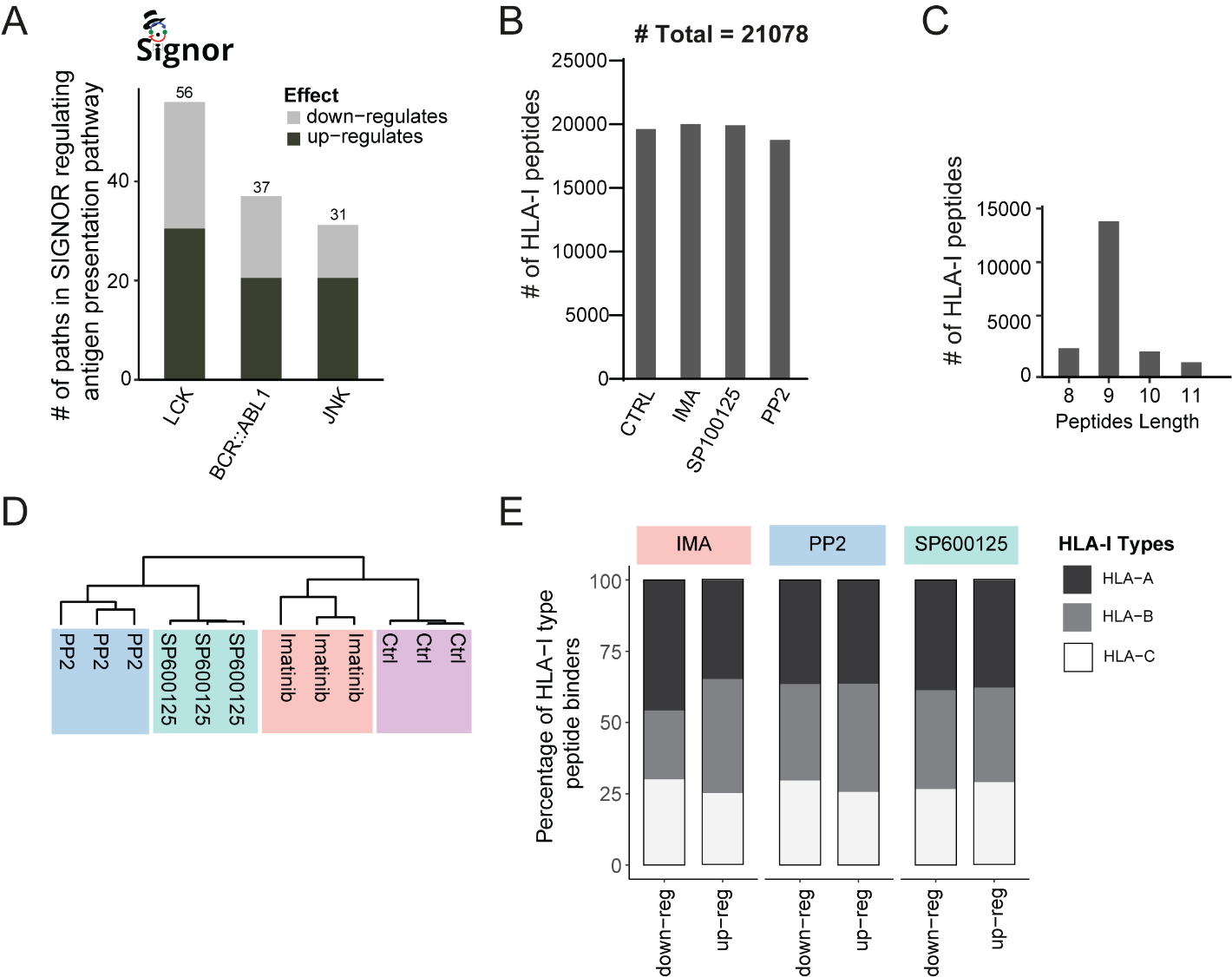

**Figure S1. A.** Barplot reporting the number of paths through which the closest kinases impact the antigen presentation pathway, as predicted by the ProxPath algorithm. **B**. Total number of HLA-I peptides identified across treatments and control. **C.** Length distribution of the identified HLA-I peptides. **D**. Unsupervised, hierarchical clustering (Pearson correlation distance) based on the abundance of 21,078 HLA-I peptides. **E**. Stacked barplots showing the percentage of significantly up- and down-regulated peptides for HLA-A, HLA-B, and HLA-C alleles. modulated HLA-I binders across HLA-A, -B, and -C alleles among up- and down-regulated peptides in cells Imatinib, PP2, and SP600125 treatments.

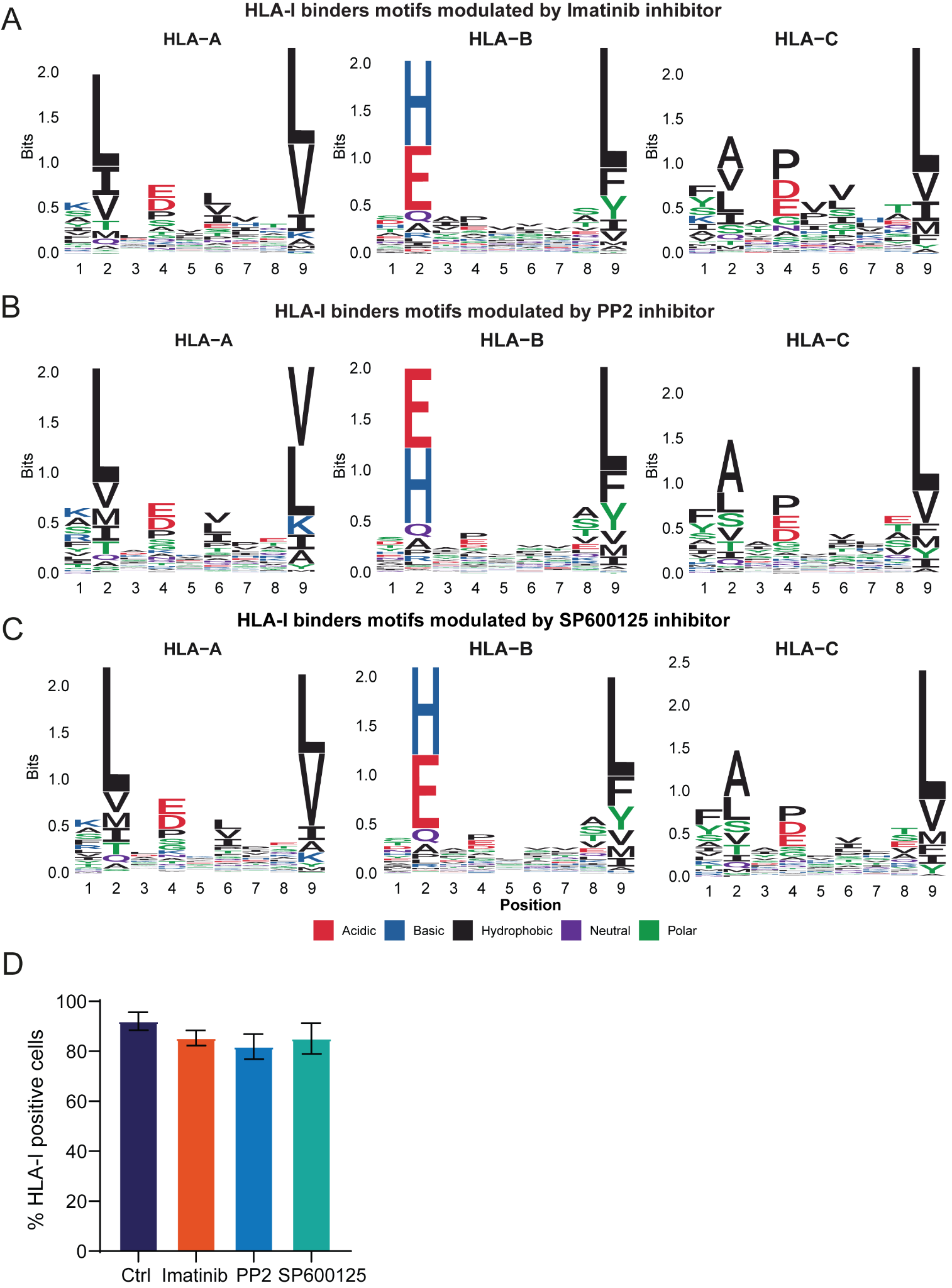

**Figure S2. A-C.** Sequence motifs of HLA-I peptides modulated by Imatinib (**A**), PP2 (**B**), and SP600125 (**C**) treatments, stratified by HLA-A, HLA-B, and HLA-C alleles. **D.** Flow cytometry analysis to assess HLA-I positive BV173 cells upon 48 hours treatment with imatinib, PP2, and SP600125.

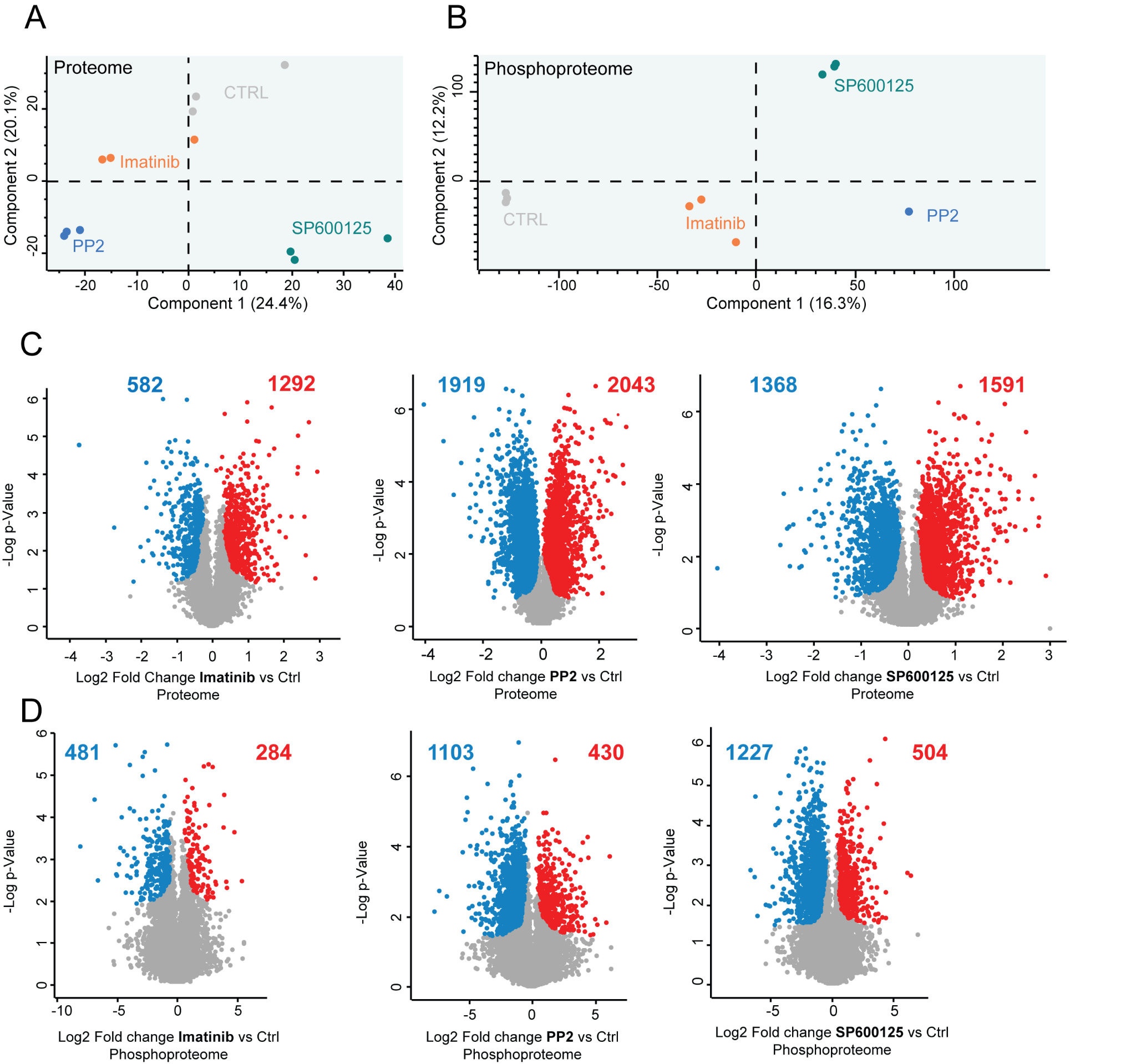

**Figure S3. A-B.** Principal component analysis (PCA) of proteins (**A**) and phosphopeptides (**B**) quantified in BV173 cells treated with three different kinase inhibitor treatments (Imatinib, PP2, SP600125) and control conditions. **C.** Volcano plots showing significantly up-regulated (red) and down-regulated (blue) proteins in imatinib, PP2, and SP600125 samples compared to control condition. Proteins that are not significantly modulated are represented in grey. how kinase inhibition impacts the exposure of novel TAAs. **D.** Volcano plots showing significantly up-regulated (red) and down-regulated (blue) proteins in imatinib, PP2, and SP600125 samples compared to control condition. Proteins that are not significantly modulated are represented in grey. how kinase inhibition impacts the exposure of novel TAAs.
